# Nanopore metagenomic sequencing of full length human metapneumovirus (HMPV) **within a unique sub-lineage**

**DOI:** 10.1101/496687

**Authors:** Yifei Xu, Kuiama Lewandowski, Sheila Lumley, Nicholas Sanderson, Ali Vaughan, Richard Vipond, Miles Carroll, Katie Jeffery, Dona Foster, A Sarah Walker, Timothy Peto, Derrick Crook, Steven T Pullan, Philippa C Matthews

**Author notes:** Address correspondence to Yifei Xu,.

## Abstract

Human metapneumovirus (HMPV) has been recognized as an important pathogen which can cause a spectrum of respiratory tract disease. Here, we report Nanopore metagenomic sequencing of the first full length HMPV genome directly from a throat swab from a UK patient with complex lung disease and immunocompromise. We found a predominance (26.4%) of HMPV reads in the metagenomic sequencing data and consequently assembled the full genome at a high depth of coverage (mean 4,786). Through phylogenetic analyses, we identified this HMPV strain to originate from a unique genetic group in A2b, showing the presence of this group in the UK. Our study demonstrated the effectiveness of Nanopore metagenomic sequencing for diagnosing infectious diseases and recovering complete sequences for genomic characterization, highlighting the applicability of Nanopore sequencing in clinical settings.

**Importance:** Nanopore metagenomic sequencing has the potential to evolve as a point-of-care test for a range of infectious diseases. Here, we report the first full length human metapneumovirus (HMPV) genome in the UK sequenced by Nanopore from a non-invasive sample from an immunocompromised patient. We demonstrate the presence of HMPV from a unique genetic group not previously reported from the UK. Our study demonstrates the effectiveness of Nanopore sequencing for diagnosing an infection that was not detected by routine first-line tests in the clinical microbiology laboratory. We report sufficient genomic data to provide insight into the epidemiology of infection and with the potential to inform treatment decisions.

## Manuscript text

Human metapneumovirus (HMPV) is a negative-sense, single-stranded RNA virus of approximately 13kb and belongs to the family Paramyxoviridae [1]. Since it was first described in 2001, HMPV has been recognized as an important pathogen which can cause respiratory tract diseases, ranging from mild upper respiratory tract infections to severe bronchiolitis and pneumonia [2]. HMPV can also cause severe disease in immunocompromised patients and those with underlying medical conditions, including lung transplant recipients [3]. Two main genetic lineages (A and B) and five sublineages (A1, A2a, A2b, B1, and B2) have been described [4].

The Nanopore sequencing platform (Oxford Nanopore Technology, ONT) is capable of generating real-time sequencing data, with the potential to evolve as a point-of-care test for a range of infectious diseases [5,6]. In this report, we describe recovery of full length HMPV genome directly from a throat swab through the application of Nanopore metagenomic sequencing.

A male in his 40’s with cystic fibrosis (CF) and a previous lung transplant presented with breathlessness, thick sputum and low oxygen saturations. His condition was further complicated by CF-related diabetes mellitus and bronchiolitis obliterans. To our knowledge, he had not travelled outside the UK. As he presented to hospital during the peak of the influenza season, a throat swab was taken to test for respiratory viruses in a clinical diagnostic laboratory; this sample was negative by PCR for influenza A, influenza B, and respiratory syncytial virus. Given his previous confirmed colonisation with *Pseudomonas aeruginosa*, he was treated with broad spectrum intravenous antibiotics, and discharged from hospital after two weeks.

We performed Nanopore metagenomic sequencing and generated 168,811 reads from this throat swab. We identified 44,580 (26.4%) HMPV reads and 5,393 (3.1%) human reads (which were discarded and not retained). The remaining reads mostly comprised bacteria representing oral flora (predominantly Lactobacilli (20%), Actinobacteria (7%), and Proteobacteria (6%)) (Fig. S1). Mapping results showed that HMPV reads covered 99.8% (13,291/13,319) of the reference sequence (USA/NM009/2016, accession number KY474539) at a high mean depth of coverage (4,786). The mean alignment length was 1,534bp and 25% of the alignments were longer than 2,000bp (Fig. 1). We used an alignment-based approach to recover a HMPV genomic sequence of 12,893bp, referred to as JR001 (accession number xxx). The sequence is nearly complete excepting 205bp at the start of the coding region.

**Figure 1.**
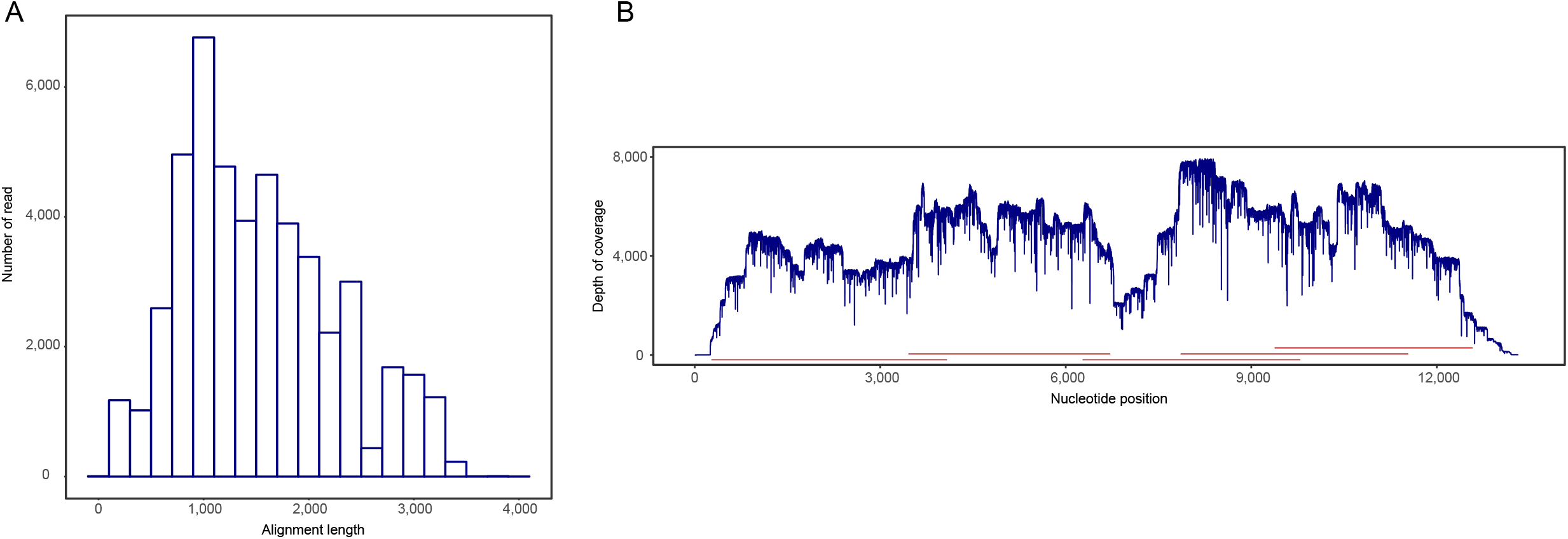
Results of Nanopore sequencing of an HMPV isolate from a throat swab. (A) Histogram of alignment length derived by mapping Nanopore reads to HMPV reference sequence (USA/NM009/2016, accession number KY474539). The mean alignment length is 1,534bp and 25% of the alignment are longer than 2,000bp. (B) Plot of depth of coverage. HMPV reads cover the full reference genome (99.8%) at a high depth of coverage (mean 4,786). Five HMPV reads, indicated by red lines, are nearly able to cover the full reference genome.

To determine the relationship between JR001 and previously published HMPV genomes, we constructed phylogenetic trees for the full length genome and eight genes (N, P, M, F, M2, SH, G, and L). JR001 clustered within genetic sublineage A2b on the basis of the full length genome and individual genes (Fig. 2 and Fig. S2). Seven HMPV strains from the United States and one strain from China were closely related to JR001, and formed a unique genetic group separated from other strains in A2b, strongly supported by a bootstrap value of 100. The pair-wise nucleotide sequence identities between JR001 and the eight related genomes ranges from 98.3% to 99.2%. This subgroup has been recently identified based on phylogenetic analysis of fusion and attachment genes [7], and comprises sequences originating from East and Southeast Asian countries, including Malaysia, Vietnam, Cambodia, China, and Japan, between 2006 and 2012 [7,8], and Croatia between 2011 and 2014 [9]. Our study provided evidence supporting the presence of HMPV from this unique group in the UK. While we found JR001 shared high nucleotide sequence identities with HMPV strains from the US, its source remains unclear. Further studies are needed to investigate the geographical distribution of this unique genetic group of HMPV and its contribution to respiratory disease in the population.

**Figure 2.**
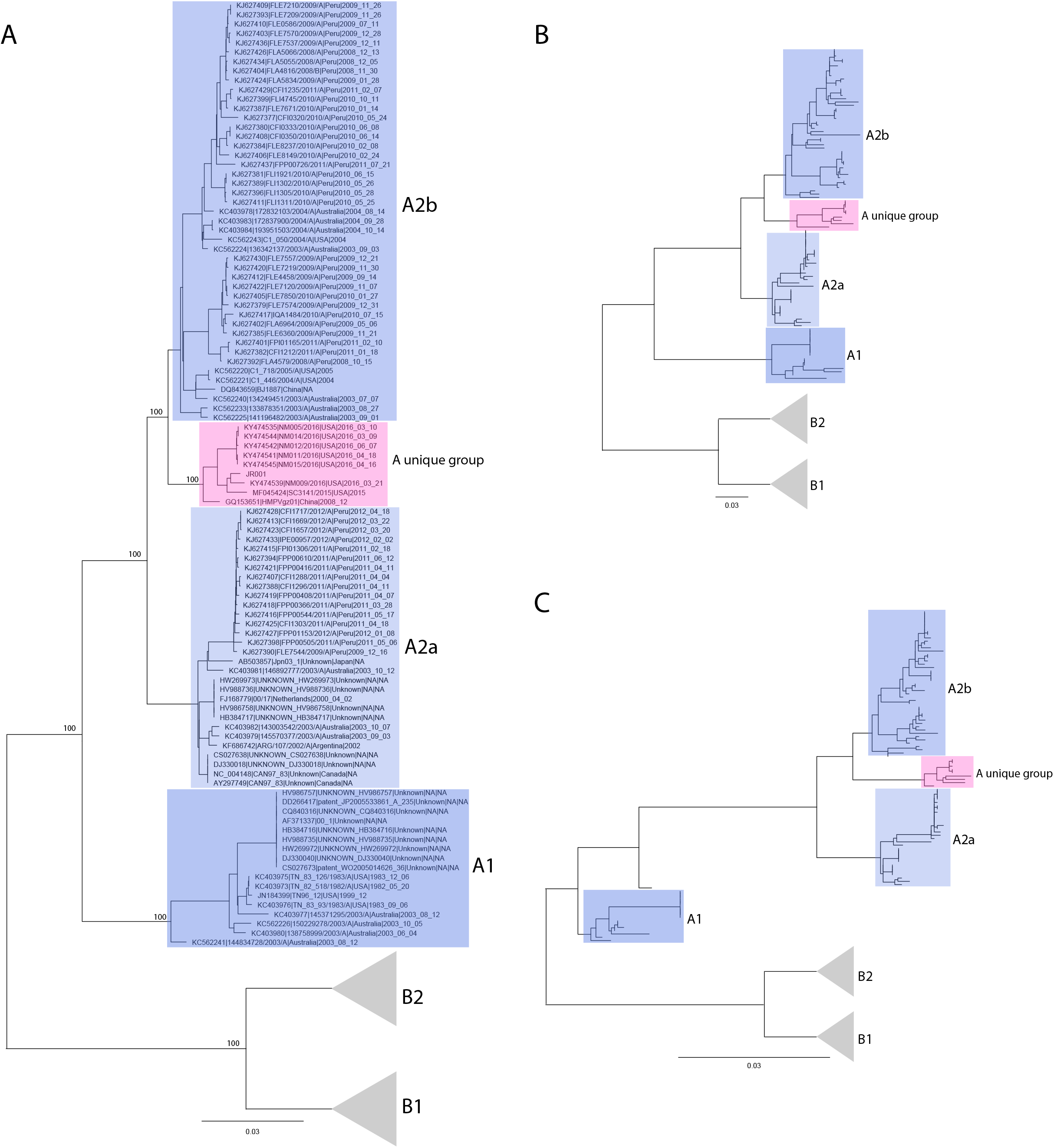
Maximum-likelihood (ML) phylogenetic trees of HMPV isolates from this study and public databases. (A) ML tree for the full HMPV genome, (B) ML tree for G gene, (C) ML tree for F gene. Five known genetic sublineages, A1, A2a, A2b, B1, and B2, are indicated by blue boxes and grey triangles. Numbers at the nodes indicate bootstrap support evaluated by 1,000 replicates. The complete phylogenies, showing name of all strains included in the analyses, are shown in supplementary Fig. S2. HMPV strain from this study and eight strains from US and China formed a unique group within A2, indicated by a red box.

We conducted time-scale phylogenetic analyses for the HMPV genome to estimate the time of emergence of this group. The topology of the time-scale phylogeny was consistent with that from the maximum-likelihood phylogenetic analyses. HMPV strains within the group were estimated to share a common ancestor originating in 2003 (95% highest posterior density [HPD], 1994 to 2008).

The extent to which the virus is a pathogen in this context is uncertain, as the patient was also at high risk of acute exacerbations of bacterial infection arising from Pseudomonas colonisation. However, the recovery of the complete genome and the predominance of HMPV reads from the metagenome suggest active infection which could have been completely or partly responsible for the acute clinical deterioration. It is not uncommon to observe co-infection of HMPV with other respiratory viral pathogens, especially respiratory syncytial virus [10]; however we did not detect sequencing reads likely to represent other significant pathogens in this case (Fig. S1).

The case is the first full length HMPV genome in the UK sequenced by Nanopore technology directly from a non-invasive sample without the need for enrichment or viral isolation, diagnosing a potentially relevant pathogen that was not detected by routine first-line tests in the clinical microbiology laboratory, and producing data that can inform treatment as well as providing insights into the epidemiology of infection. Characterisation of the microbiome of patients with complex underlying lung disease, both during periods of clinical stability and in the setting of lower respiratory tract infections, could be valuable in informing intervention and supporting antimicrobial stewardship.

## Methods

### Sample collection, preparation, and Nanopore sequencing

A throat swab was collected in viral transport media from a patient presenting to our tertiary referral teaching hospital in Oxford, United Kingdom. The sample was tested for respiratory viruses using Xpert Xpress Flu/RSV assay (Cepheid, Sunnyvale, CA, USA) in a clinical diagnostic laboratory. The sample was frozen for retrospective Nanopore sequencing. The sample was thawed and passed through a 0.45 μm filter prior to RNA extraction and DNase treatment. cDNA was prepared and amplified using a Sequence-Independent-Single-Primer-Amplification method as described previously [11]. cDNA was used as input for a SQK-LS108 library preparation and sequencing on a R9.4.1 flow cell using a MinION device (ONT).

### Genomic analysis

Nanopore reads were basecalled using Albacore v2.1.7 (ONT). Metagenomic classification and mapping were used to identify HMPV reads. Reads were first taxonomically classified against RefSeq database using Centrifuge v1.0.3 [12]. *De novo* assembly was then performed with HMPV-like reads using Canu v1.7 [13]. The resulting contigs were BLASTed against GenBank nt database to determine the reference HMPV sequence. Reads were mapped against the selected reference (USA/NM009/2016, accession number KY474539) using Minimap2 [14]. HMPV reads were defined as those assigned to HMPV by centrifuge and confirmed by mapping. Consensus sequence for the HMPV strain was built using Nanopolish v0.9.2 [15].

Phylogenetic analyses were conducted using an integrated dataset that comprised the HMPV sequence from this study and 154 complete HMPV genomic sequences from NIAID Virus Pathogen Database and Analysis Resource (ViPR) and NCBI Genbank [16]. Maximum-likelihood phylogenies were generated using RAxML v8.2.10 [17]. Time scale phylogenies were built for genomic sequences with complete sampling dates (month, day, and year) using BEAST v1.10.1 [18]. The SRD06 partitioned substitution model, uncorrelated lognormal relaxed clock model, and Bayesian skyline coalescent tree prior were used in the analyses. Multiple independent runs were performed with a chain length of 200 million steps and sampled every 10,000 steps. These runs were combined to ensure an adequate effective sample size (>200) for relevant parameters.

## Ethics statement

This sample that was surplus to diagnostic requirements was sequenced as part of a larger study with Research Ethics Committee approval (17/L0/1420).

## Accession number

The sequencing data was deposited in the xxx under accession no. xxx.

## Funding

This work was supported by NIHR Oxford Biomedical Research Centre.

## Supplemental materials

**Figure S1.** Taxonomic assignment of Nanopore sequencing reads of a throat swab from a patient with complex lung disease and immunocompromise. HMPV reads accounted for 26% of the total reads.

**Figure S2.** Maximum-likelihood (ML) phylogenetic trees for the full length genome and gene of HMPV isolates from this study and public databases. Numbers at the nodes indicate bootstrap support evaluated by 1,000 replicates. Five known genetic sublineages, A1, A2a, A2b, B1, and B2, are indicated by blue and grey boxes. HMPV strain from this study and eight strains from US and China formed a unique group within A2, indicated by a red box.

